# A dynamic causal inference framework for perception–action loops

**DOI:** 10.1101/2025.11.13.688198

**Authors:** James R.H. Cooke, W. Pieter Medendorp

## Abstract

Causal inference, the process of inferring the causes of our sensory input, is central to multisensory perception. While most computational models of causal inference focus on static perceptual tasks with no temporal or motor components, real-world behavior unfolds dynamically and often involves closed-loop control. Here, we introduce and validate a general modeling framework that unifies multisensory causal inference with optimal feedback control by casting the problem as inference and action in a switching linear dynamical system (SLDS). Our framework combines approximate inference over latent causal structures with a mixture-of-controllers approach, in which each controller is optimized for a specific causal model and weighted by the current belief in that model. We show that classical static models of multisensory perception are special cases of our framework, and extend them to dynamic, action-oriented settings where inference and control evolve over time. Using simulations of previously published behavioral tasks, including visuo-vestibular heading discrimination and path integration with interception, we show that the model reproduces key empirical findings, such as S-shaped biases in heading estimation and motor strategies based on inferred object motion. We also demonstrate the versatility of our approach in novel task extensions, including motion in depth, latent switching dynamics, and multi-objective motor control. Together, our results provide a principled, general-purpose computational account of how causal inference and motor behavior interact in dynamic, uncertain environments, offering a bridge between theories of perception and action.

**Author summary:** When we interact with the world, our brain constantly combines information from different senses — like visual and vestibular sensory input - to determine what is happening and guide an appropriate response. However, these signals do not always originate from the same source in the world. For example, a moving object might not match the background movement we feel during self-motion. In such situations, the brain faces the challenge of “causal inference”: do these signal arise from a common source or from separate events.

Most previous research has studied this problem in simplified, perceptual settings. But in everyday life, we move through the world and make decisions in real time based on noisy and often conflicting sensory input. In this work, we develop and validate a new computational framework that combines causal inference with models of movement control. This approach allows us to simulate more realistic situations where perception and action interact continuously. We show that our model not only reproduces results from earlier experiments but also makes predictions about how people behave in more complex tasks, like intercepting moving targets or navigating uncertain environments. Our framework offers a new way to study how the brain integrates sensory information to guide behavior in the real world.

## Introduction

In our daily lives, we navigate complex environments without direct access to the true states of objects and surroundings. Instead, our sensory systems provide noisy and incomplete information [1, 2], which introduces uncertainty in perception. This raises a fundamental question: how can the brain effectively use imperfect sensory cues to accurately infer the state of the environment?

Cues from the same sensory modality, such as visual depth and texture indicating object slant, typically provide consistent information. Over time, the brain learns to integrate these cues optimally, resulting in a more precise representation of the tilt of the object [3, 4]. Similarly, combining cues across different sensory modalities, such as visual and vestibular inputs, can improve the precision of estimates of spatial orientation and self-motion [5].

However, indiscriminate fusion of cues from multiple modalities can lead to errors when the signals originate from different sources or represent different states of the world. In self-motion estimation, for instance, optic flow detected by the visual system may stem from the observer’s own movement, from independently moving objects [6], or a mixture of both. The brain must disentangle these contributions and integrate the relevant component with inertial motion cues when available [7].

Thus, correctly inferring the state of the environment (whether we, the object, or both are moving) depends on deciding whether sensory cues should be integrated or kept segregated. Conceptually, this represents a hierarchical inference problem, involving both estimating latent environmental states and inferring the causal structure that links sensory signals.

Bayesian causal inference (BCI) frameworks formalize this hierarchical inference by explicitly representing both the probabilistic nature of sensory information and the underlying causal structure [8, 9]. Empirical evidence supports this view, showing that distinct brain areas represent different stages of BCI computations, including unisensory processing, multisensory integration, and causal inference [10].

Although BCI effectively accounts for many aspects of perceptual inference, the experimental paradigms used to support it typically rely on open-loop, trial-based psychometric tasks requiring discrete judgements [7, 8, 11]. Even in studies using dynamic stimuli, the time-varying sequential sensory inputs are typically reduced to aggregated observations, leaving the framework’s applicability to continuous sensorimotor behaviors largely untested [7, 11]. Indeed, such reductions are inadequate when considering real-world tasks, like cycling or driving, that require ongoing, closed-loop integration of sensory processing and motor control.

In the field of human motor control it is thought that the brain estimates the state of the world and body using a combination of sensory and motor feedback and then uses this state to generate appropriate actions. A prominent probabilistic framework capturing this idea is optimal feedback control (OFC), often formalized within a linear-quadratic-Gaussian (LQG) setting [12–15]. OFC posits that motor behavior arises from an optimal controller minimizing a cost function (e.g., effort, accuracy) in tandem with an optimal state estimator that integrates sensory feedback based on a generative model. However, hitherto OFC has not accounted for dynamic perceptual causal inference - the ability to handle uncertainty about the causal structure linking sensory variables which can vary over time.

To provide a unified computational account of sensorimotor behavior, it is necessary to integrate Bayesian causal inference with the continuous control optimization of OFC. Here, we address this gap by developing a framework that jointly performs causal inference and continuous motor control under uncertainty. We apply this framework to the problem of steering using optic flow in dynamic environments and demonstrate that it can capture a broader array of adaptive sensorimotor behavior than existing approaches.

## Results

### Modelling framework

Much of the work on causal inference in perception has focused on tasks which can be effectively summarized via the final time point of the task [7, 8, 11, 16–18].For instance, Acerbi et al. [11] performed a rigorous model comparison to understand how observers estimate heading direction in the presence of conflicting visual and vestibular cues. Their approach involved collapsing the time-varying sensory inputs into single visual and vestibular measurements of heading. While effective for their research aims, such an approach precludes modelling how sensory information is processed over time, thereby limiting insights into the interaction between the continuous evolution of causal inference and closed-loop control.

In the following, we first show that that previous casual inference models in multisensory perception can be understood as sub-cases within our framework [7, 8, 11, 16, 17]. This perspective allows us to build our framework and embedded algorithm up progressively, starting from the simplest static case of causal inference (two measurements, no control or dynamics), extending this to dynamic causal inference across time, and finally integrating closed loop control (see Fig 1A).

**Fig 1.**
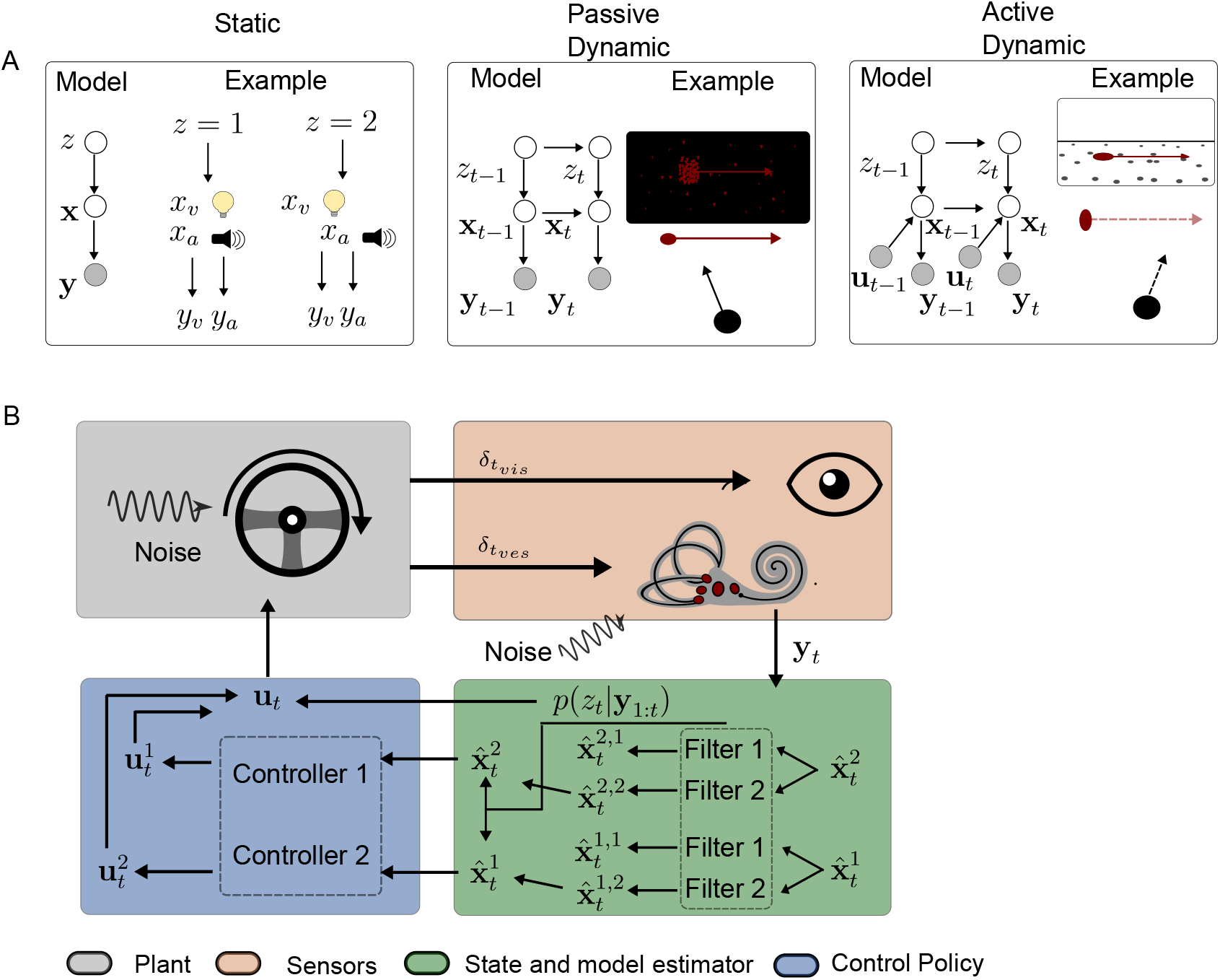
Categories of causal inference problems alongside a unified algorithm to handle them. A. Classification of causal inference tasks, from the simplest static case to the most complex case involving closed-loop control. For each case, on the left, the corresponding generative model from the observer’s standpoint (in gray, observed variables; white, unobserved), and on the right, a specific example. Variables denoted **x** indicates relevant state variables, *z* the causal structure and **y** the observations. Examples indicate a simple audio-visual task (static) [8], a heading based task in the presence of object motion (passive dynamic) [7], and a target interception task when the target moves on some trials but not on others [19] (active dynamic). B. Schematic overview of our framework, applied to the active dynamic task or similar tasks [20, 21]. The observer must steer to land on target. The visual and vestibular system provide noisy and delayed measurements of both the target and self-motion (orange box). These measurements are used to compute state estimates (target, motion), conditioned on the probability of each causal structure (green box). An optimal motor command is computed for each model, and weighted by the corresponding probability, to give a final motor command to turn the wheel (blue box).

Doing so yields a novel framework that combines elements of optimal feedback control [12, 22, 23] with causal inference over time [19]. Specifically, we formalize multisensory causal inference as performing inference on a switching linear dynamical system (SLDS) [24]. In this type of system, the structure (matrices) of the system change over time (switches), so solving for the optimal motor command in such a system is intractable. To handle this, we propose a heuristic approach based on mixture of sub-controllers, each optimized for a distinct causal model [25]. Fig 1B indicates a schematic of how the algorithm works for a naturalistic style steering task (similar to Fig 1A, active dynamic).

### Problem specification

We consider a control problem in which a participant must act based on uncertain sensory information while also inferring the causal structure that generated these observations. Specifically, we consider a system which switches between separate linear Gaussian systems [24] (SLDS).

Formally, at each time step *t*, the true causal model is indicated by a latent variable *z*_*t*_ ∈ {1, …, *M*}, which evolves according to a Markov process with known transition probabilities

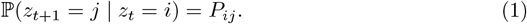

Each model ℳ_*j*_ defines a distinct linear-Gaussian system with latent state 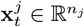, dynamics:

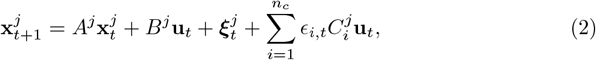

and observation model:

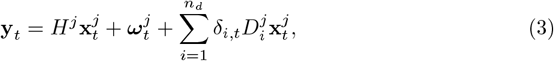

where **u**_*t*_ ∈ ℝ^*m*^ is the control signal. The noise terms 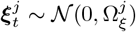 and 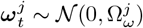 represent additive Gaussian noise, while *ϵ*_*i,t*_, *δ*_*i,t*_ ~ 𝒩 (0, 1) denote signal-dependent (multiplicative) noise.

### Static CI

We can view the majority of multisensory causal inference models as static, single-step examples [7, 8, 16, 17] of the system defined by Eq 1, 2, and 3. In these models, the observer receives a single set of multisensory measurements, typically, one from each modality (e.g., auditory and visual cues), and must both infer the underlying causal structure and estimate the latent stimulus variable (e.g., visual location or heading direction). The full marginal likelihood of the observation is given by a mixture of Gaussians, so posterior inference over *z* and **x** can then be computed via Bayes rule (see [26]). The observer’s response is then typically modelled as the estimate that minimizes expected loss under the inferred posterior [8, 11], For example, the observer may respond with the mean of the posterior (which minimizes the squared error with the true stimulus). Under these assumptions, classic static causal inference models can be understood as a special case of a more general switching state-space model, simplified by the absence of temporal dynamics: *A* = 0, *B* = 0, *T* = 1 and no control inputs: **u**_*t*_ = 0.

### Example: Audio-Visual Localization

To illustrate this more concretely, consider the canonical example of static causal inference: audio-visual localization [8] (see Fig 1.A, Static). In this task, an observer is presented with simultaneous auditory and visual cues originating either from the same location or from different locations and is asked to judge whether the cues arise from a common source, or to estimate the location of one of the cues.

For modelling, the original data (e.g., auditory time series, multiple visual frames) are reduced to scalar measurements of location: one visual (*x*_*v*_) and one auditory (*x*_*a*_). The observer receives noisy observations of these latent locations:

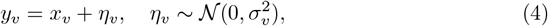

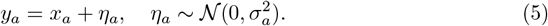

The latent locations can arise under two causal structures, *c* = 1: Common source — *x*_*v*_ = *x*_*a*_ = *x*, drawn from a prior 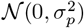 or *c* = 2: Independent sources — *x*_*v*_, *x*_*a*_ drawn independently from 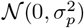.

This can be expressed as a one-step switching linear-Gaussian model. Under the common-source model (*c* = 1), we can define:

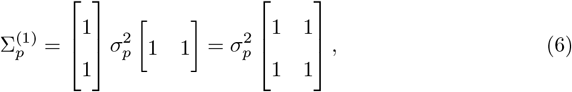

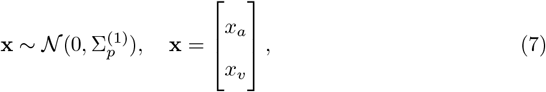

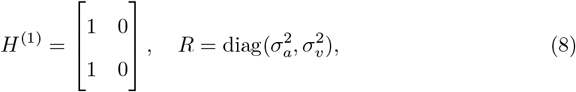

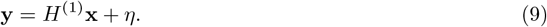

where *x* is the concatenated state, *R* indicates the sensory noise matrix (for auditory and visual measurements), Σ_*p*_ indicates the a prior state covariance, *H* is the measurement matrix and *η* is a zero-mean normally distributed noise with covariance *R*.

Under the independent-source model (*c* = 2), we assume uncorrelated latent sources:

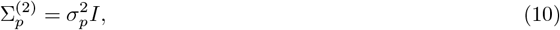

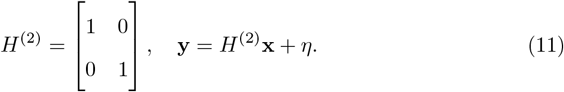

By modelling this static problem as a linear system, we establish a conceptual and mathematical bridge between earlier work on perceptual causal inference work and dynamic sensorimotor control. However, in order to model more complex and naturalistic tasks, this static model must be extended to a temporally unfolding, closed-loop control setting, where both inference and action co-evolve over time.

### Passive dynamic CI

We begin by generalizing from static causal inference to dynamic causal inference. This means the participant now receives a stream of sensory inputs over time and must use these inputs to estimate the time-varying environmental state and estimate the underlying causal model of the sensory observations (see Fig 1A, Passive dynamic).

This amounts to a switching linear system but without any controlled input (*B* = 0), so we must specify the dynamics (*A* matrix) and observational characteristics (*H*) of the different models as well as the noise terms. Unfortunately, this transition from static to dynamic makes the inference no longer tractable. This is due to the number of potential mode sequences *z*_1:*T*_ increasing exponentially over time. As such, it is necessary to resort to some form of approximate inference to handle combined state estimation and causal inference. We opt for using the Generalized Pseudobayesian Estimator of order 2 [24]. The basic idea behind this approach is approximate this intractable mixture of linear systems with a finite mixture of linear systems. This allows us model the system with an approximate probability of the different causal structures (the weighting of the Gaussians) alongside a state estimate and corresponding covariance of the state estimate. These state estimate are obtained by considering the potential state estimates that could result from different models, and weighting these state estimates by the likelihood of the different models (see Methods for details).

#### Example : Visuo-vestibular heading inference over time

As an illustrative case, we consider a recently published visual-vestibular causal inference task, in which the observer must decide if an object is stationary or moving during concurrent self-motion [7, 17]. In this task, participants were passively moved and presented with optic flow consistent with this motion, except for regions containing an independent object that could be either stationary or moving at various speeds.

Participant were asked to judge whether their heading direction deviated from straight ahead and report whether the independent object was stationary or in motion. To model this experiment, the author’s summarized a full trial via a scalar measurement of the object’s velocity, the background flow velocity, and the participant’s inertial motion (as sensed by the vestibular system). With our framework, we model the observer’s inference process over time, capturing the evolving estimate of self-motion and object motion as well as the associated model probabilities. To develop this dynamic case, we must explicitly define the observer’s internal beliefs about the dynamics of their own body and of the object. To this end (see Methods), we model the body as a point mass along a minimum jerk trajectory with some heading angle, while the independent object moves with a Gaussian velocity profile with different peak speeds. Figure 2 presents the simulation results, alongside the original experimental data.

**Fig 2.**
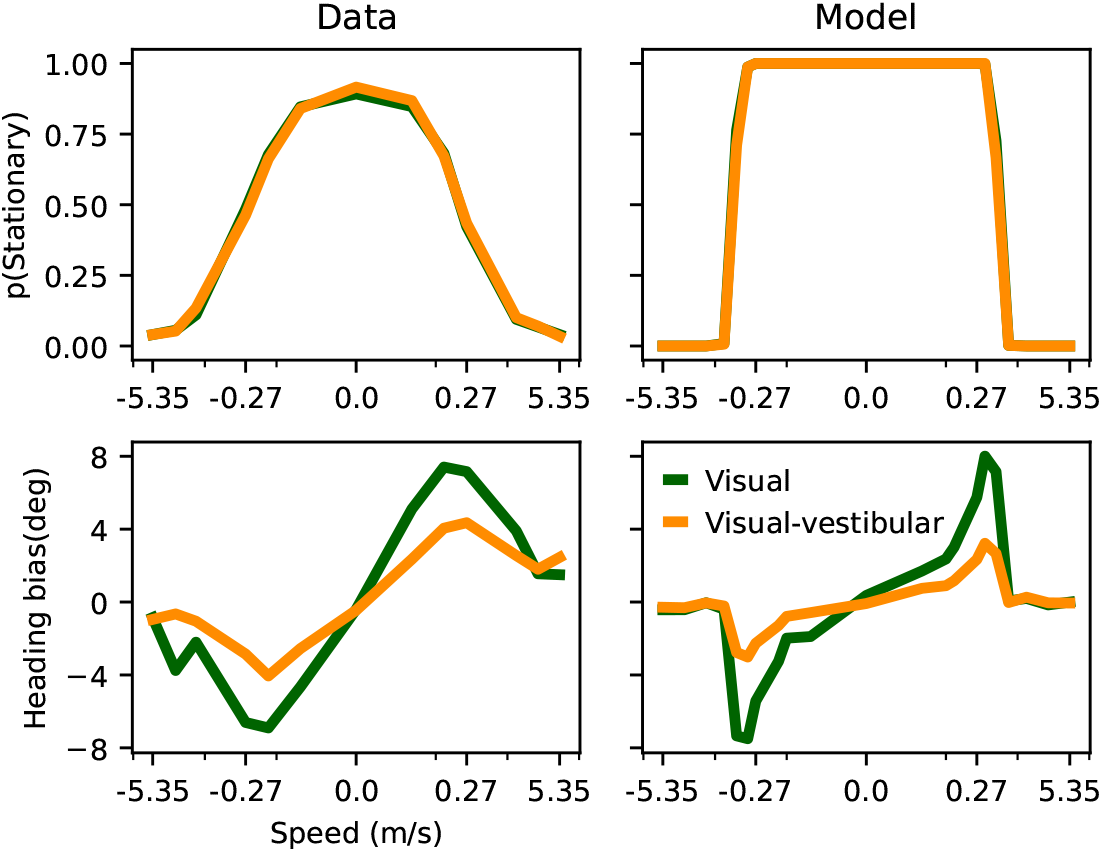
Data (left) and model predictions (right) for a visual-vestibular causal inference task. Top: detection of independent object motion as a function of object speed. Bottom: heading bias as a function of the independent object speed. Data from [7].

The model successfully captures the qualitative patterns observed in the data. For stationary perception (Fig 2, top panels), slower object motion increases the likelihood that observers perceive it as stationary, with little difference across modality conditions. For heading bias (Fig 2, bottom panels), the simulations reproduced the characteristics ‘S-shaped’ pattern: at low speeds, participants integrate the object information with their own estimate of self-motion, producing a bias: as speed increases the object information is weighted less and so the bias decreases again. Both the empirical data and the model simulations show a more pronounced effect under vision-only conditions (as the vestibular system provides an additional measurement unaffected by the object). Thus, we can relax many of the assumptions of previous causal inference models [7, 8, 16] while predicting the observed pattern of results.

### Active dynamic CI

Up to this point, we have considered causal inference in both static and dynamic perceptual settings, where the observer passively receives sensory input. However, in real-world behavior, observers are active participants that continuously generate motor commands based on their internal beliefs. This introduces an additional layer of complexity: actions shape future sensory input and thus alter the process of causal inference itself. Addressing this challenge requires a unified framework that combines causal inference with closed-loop control. From the participant’s perspective, two tasks must be performed simultaneously: estimate the current state 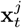 and causal model while selecting a control **u**_*t*_ input that drive the system toward accomplishing the task.

Because exact inference and control in SLDS is intractable, we combined an approximate inference method - the Generalized Pseudo Bayesian algorithm of order 2 (GPB2) [24]-with a mixture of sub-controllers [25], where each controller is designed to be approximately optimal for a particular causal model (see Methods for details of the algorithm). The final control signal is a weighted mixture

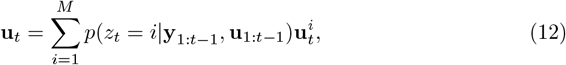

where 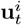 is the control computed under model ℳ_*i*_, and *p*(*z*_*t*_ = *i*|**y**_1:*t*−1_, **u**_1:*t*−1_) is the approximate posterior belief over the causal structure (see Methods). Each controller minimizes a cost function of the form (for ℳ_*j*_)

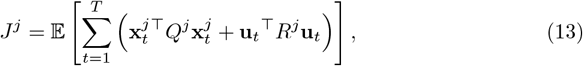

subject to the dynamics of model ℳ_*j*_, the state cost matrix *Q*^*j*^ and control cost matrix *R*^*j*^.

This framework allows the participant to behave in a way that is robust to uncertainty about causal structure, continuously adjusting its control policy as its beliefs evolve over time. Crucially, the causal inference process itself becomes dynamic, beliefs over *z*_*t*_ influence action, which in turn shape future observations and thus future beliefs.

#### Example: Path integration with interception

As an illustration, we modeled the path integration interception task used by Noel et al [19].In this task, participants were briefly (75 or 150 ms) presented with a target at a fixed depth but varying direction, which either remained stationary or moved laterally at a constant speed. Using a joystick, participants then navigated to intercept the target. Their steering endpoints were analyzed to assess how accurately they had estimated the target’s speed. At the end of each trial, participants also reported whether they perceived the target as moving or stationary. The authors modeled their data by assuming perfect knowledge of target depth and relying on an open-loop interception policy. Our general framework, based on switching linear dynamical systems with feedback control, removes these assumptions and requires depth to be estimated and incorporates a closed-loop control for the interception.

The resulting predictions are shown alongside the data in Fig 3. Fig 3 displays the proportion of trials in which participants (top panels) and the model (bottom panels) judged the target as stationary, plotted against target speed (x-axis). Results are separated by whether the participant was moving during target presentation (self-motion, red) or stationary (black), and by target presentation time (left vs. right panels). The bottom four panels show the estimated target speed, based on the steering endpoints, in the same format. As shown, the model predictions align well with the qualitative trends in the data. Accuracy in both stationary reports and estimated speed becomes more accurate as the amount of available data (e.g. presentation time) increases, particularly when no self-motion is present. In addition, the characteristic ‘S-shaped’ pattern emerged in estimated speed: participants show stronger integration at low target speeds, inducing a bias, but shift to segregation at higher speeds, resulting in more veridical speed estimates.

**Fig 3.**
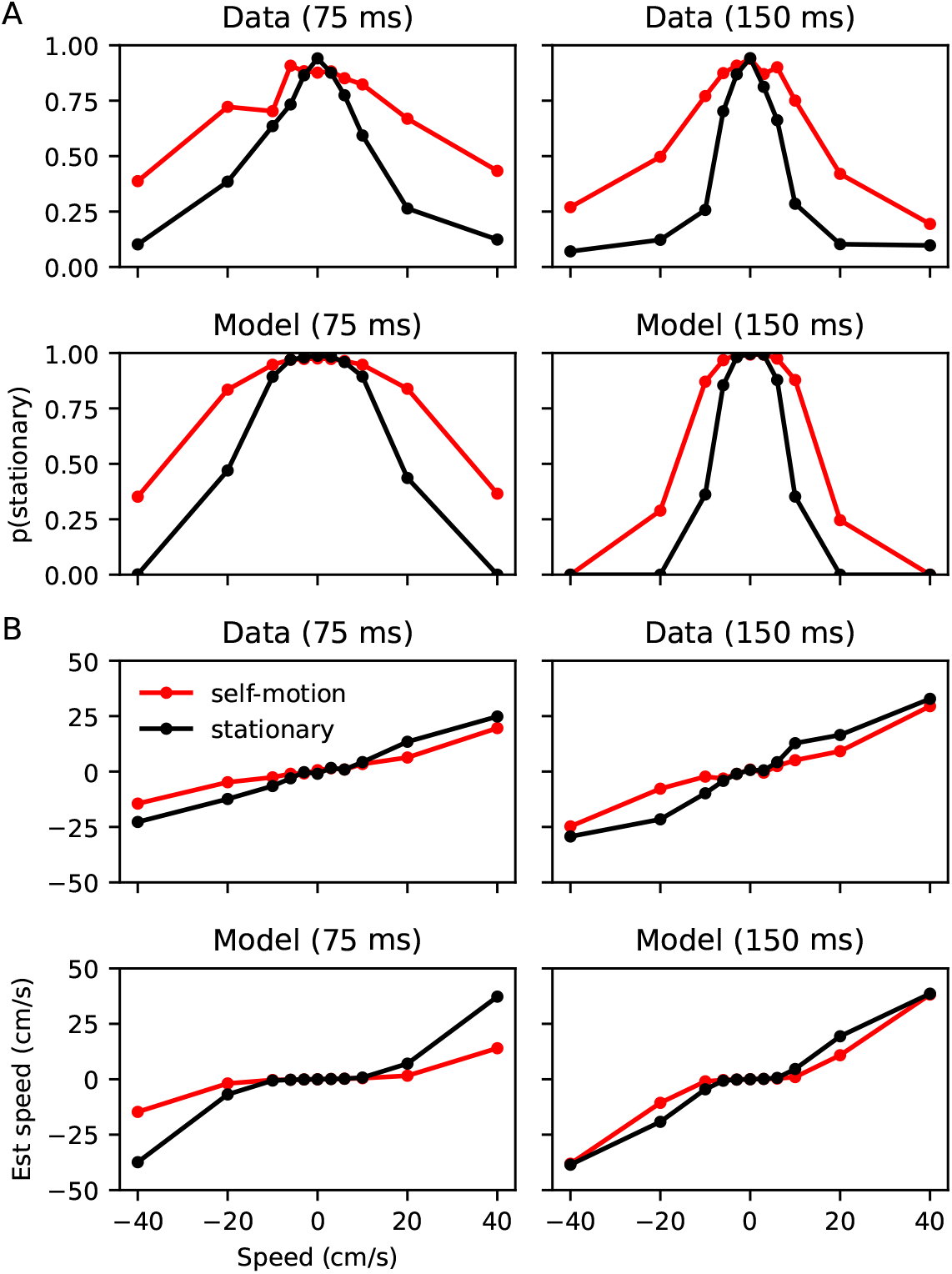
Data alongside model predictions for Noel et al. task [19]. A. The top four panels show the proportion of trials in which participants (top row) and the model (bottom rows) judged the target as stationary, plotted against target speed (x-axis). Results are separated by participant self-motion during target presentation (red: self-motion, black: stationary) and by target presentation time (left vs. right panels). B. Estimated target speed, derived from steering endpoints, in the same format.

### Prediction of the SLDS framework for more complex tasks

The model simulations so far show our framework yields predictions consistent with previous experimental findings. Importantly, our approach makes it fairly straightforward to construct participant models for more complex tasks with fewer necessary restrictions. To demonstrate this, we will examine three extensions of the interception task: (i) interception while the target moves in depth, (ii) interception when the target’s motion changes over time, and (iii) interception while passing through a via point.

#### i) Interception while the target is moving in depth

Relaxing the assumption of a known target depth in our model, a natural follow-up is to consider an interception task where the target can move in depth (see Fig 4A, top). The predicted average trajectories reveal comparable patterns to for depth-fixed interception, with the characteristic S-shaped curve for the endpoint error (Fig 4A, middle panels), as well as the inverted U-shape for the stationary reports (Fig 4A, bottom panel).

**Fig 4.**
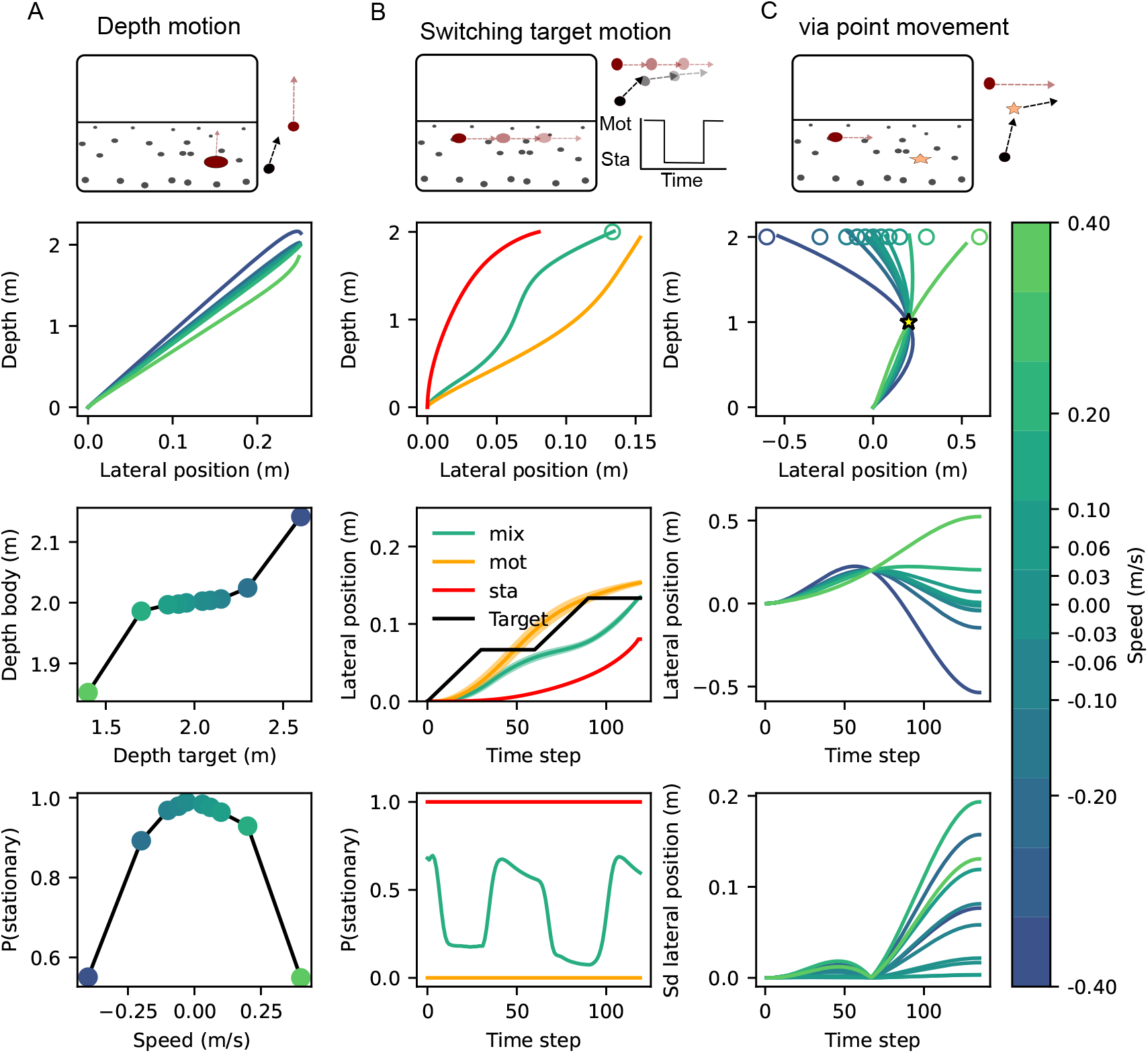
Schematic of task extensions alongside model predictions. A. Target moving in depth. Top: cartoon-like illustration of the task. Middle panels: average body trajectories across all tested speeds and endpoint error as a function of target depth. Bottom: perceived speed relative to stationary judgements. B. Changing target motion. Top: task schematic. Middle panels: average trajectories for the switching (SLDS) model (green) and non-switching models (stationary (red) and motion (orange)); only the SLDS model reaches the target. Bottom: time-varying probabilities assigned by the SLDS model to different target motion modes. C. Passing through a via point. Top panel, task schematic. Middle panels: SLDS predictions showing that movements naturally pass through the via point en route to the target. Bottom: movement variability decreases near the via point and increases again toward the target.

#### ii) Interception while the changing target motion changes over time

In the real world, objects rarely move at a constant speed. Instead, they speed up or slow down as time passes. Our framework accounts for this by having multiple motion models alongside a non-identity transition matrix. To simulate such behavior, we extend the interception task by introducing a latent switching dynamic that alternates the target being stationary and in motion (Fig 4B, top panel) Specifically, at each time step *t*, the target’s state *z*_*t*_ ∈ {0, 1} follows a deterministic periodic schedule:

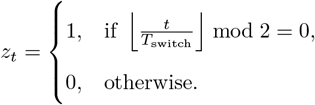

When *z*_*t*_ = 1, the target moves at a constant speed along the lateral (x) axis, while it remains stationary when *z*_*t*_ = 0. Furthermore, the target remains fully visible for the entire trial and its depth is held constant throughout.

From the average trajectories (Fig 4), we can observe distinct predictions between a switching (SLDS) model and the non-switching models (stationary only (red) or motion only (in orange)), with the switching model showing an ‘S’ like curve as the motion switches over time and reaches the target (Fig 4B, third panel). In the model, this occurs due to a change in the probability of the different modes inferred by our algorithm (Fig 4B, middle row, bottom panel) as the target motion changes over time. By contrast the non-switching models display significant error through the movement and do not end up on target (fig 4B, middle panels).

#### iii) Interception while passing through a via point

Naturalistic motor control often involves multiple task requirements (e.g., during driving we want to reach a location while adhering to road rules). At the same time, there is uncertainty in the underlying model of the world (e.g., the physics of the car and motion of other cars and pedestrians). Our framework accounts for both simultaneously. Indeed, the relative importance and structure of the task requirements significantly alters the optimal control policy [12].

To illustrate this, we examine the interception task with the additional constraint that the participant must pass through a designated via-point halfway through the movement time (*t*_via_ = *T/*2). In the model, this is implemented by augmenting the cost state (and cost function) with an additional time-specific penalty incorporated in to the Q term,

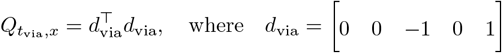

This additional cost term promotes alignment between the participant’s position and the via-point in spatial coordinates at the specified time. Importantly, the dynamics, sensory observations, and control structure remain unchanged. This illustrates a key advantage of the linear-quadratic-Gaussian framework: complex motor behaviours can be flexibly shaped by modifying the objective function rather than by altering the environment or system dynamics.

Fig 4C shows the prediction of our model. From the average trajectories (Fig 4, middle panels), we can observe that the movements naturally pass through the via point on their way towards the target rather than aiming to hit the via point first and then redirecting to the target. Fig 4, bottom panel depicts the standard deviation of the body position, showing that it decreases significantly near the via point and then increases again towards the target.

## Discussion

Here we introduced a unified framework for joint perceptual causal inference and closed-loop control, tailored toward biological systems. We formulated the problem as one of inference and control within a switching linear dynamical system (SLDS) [24] with multiplicative control and observational noise [13]. This SLDS formulation enabled the use of the efficient GPB2 algorithm [24] for inference of the causal model while also accounting for additive and multiplicative noise in state estimation and control. Our approach makes it straightforward to build causal inference-based models for tasks more complex than those previously studied. We demonstrated this by extending a static CI model [7] into a dynamic version and showed it predicts the data equally well, while relaxing previous restrictions on control-based CI tasks. This allows our framework to handle more realistic task scenarios, extending it to novel stimulus dimensions, stimulus motion profiles and additional task requirements.

Although we did not explicitly model delays, our SLDS makes this straightforward through system augmentation [27]. Such an extension would be crucial for accurately integrating information over time, ensuring that sensory signals are not only weighted by their reliability (when they represent common variables) but also by their respective delays within each sensory loop.

### Relationship to previous work

While our SLDS approach was designed to bridge the gap between static causal inference and closed-loop control, it shares conceptual parallels with established models in the motor control literature. In particular, our approach aligns closely with the MOSAIC framework [28] in that both maintain multiple internal models and combine their outputs to generate control signals. However, a key difference lies in the treatment of model knowledge and learning. MOSAIC primarily focuses on online learning of both the forward (predictive) and inverse (control) models from sensorimotor experience, adapting responsibility weights based on prediction errors to mediate model switching and learning. In contrast, our framework assumes a known set of candidate models with fixed parameters and focuses on performing inference and control under causal model uncertainty. Instead of error-driven responsibility weights, we use a principled Bayesian inference approach (GPB2) over a Markov-switching latent variable to update beliefs about the active model. This allows us to integrate causal inference and control while accounting explicitly for additive and multiplicative noise, model switching dynamics, and differing internal model structures. It also allows us to change the task requirements (e.g., a via point) straightforwardly and compute a new optimal policy. Thus, while MOSAIC emphasizes adaptive model acquisition and learning, our SLDS approach emphasizes inference and control with uncertainty over a predefined set of models.

Furthermore, our framework differs from the recently proposed COIN model of contextual inference [29], which also addresses uncertainty regarding latent generative structures. While both frameworks involve inference over a hidden switching process, they diverge in purpose and scope. COIN is primarily designed to explain behavioral adaptation patterns by inferring discrete latent contexts that govern learning rates and memory recall, without modeling real-time control or system dynamics. In contrast, our SLDS framework integrates causal inference with continuous control by modeling uncertainty over latent dynamical systems and computing model-specific control policies. Moreover, our model can explicitly handle sensorimotor features such as sensory delays and signal-dependent noise — features that are central to biological control but are not incorporated in the COIN model. Thus, while both models deal with uncertainty over internal models, our approach addresses the closed-loop control problem in continuous time under dynamically switching latent causes.

In our SLDS framework, we chose to compute the control signal as a weighted average of the control signals from each candidate model. This choice parallels Bayesian model averaging, as commonly employed in many causal inference models [7], where the final perceptual or motor estimate integrates across uncertainty in latent causes. It is important to note, though, that is only one of the plausible decision strategies. An alternative is posterior probability matching [30], which suggests that the participant samples a causal model in proportion to its posterior probability and then selects the optimal action conditioned on that sampled model. This sampling-based approach has natural parallels to Thompson sampling in decision making under uncertainty, where a participant samples a hypothesis from the posterior and acts optimally with respect to it [31, 32]. If the causal model remains fixed over time, this equivalence becomes exact. This connection underscores a deeper conceptual link between perceptual causal inference and frameworks for decision making and control under uncertainty, including those based on Thompson sampling and Bayesian reinforcement learning [33]. More generally, these strategies can be understood within the framework of bounded rationality and information-theoretic decision-making [34], which formalizes the trade-off between maximizing expected reward and minimizing information-processing costs. From this perspective, averaging control signals corresponds to a deterministic, Bayes-optimal policy, while sampling-based approaches represent resource-efficient approximations that introduce stochasticity, exploration, and potentially biologically plausible mechanisms.

### Future directions

We could improve the realism of the simulations presented here by including other aspects of sensorimotor control, such as sensorimotor delays, which may vary across modalities [35], as well as state and control-dependent noise [36].Our framework is specifically designed to integrate these factors into the optimal policies needed for control.

One key limitation of the current approach is its reliance on linear Gaussian dynamics, which enables exact inference via Kalman filtering. A natural next step is to consider non-linear, non-Gaussian systems, which are more representative of real-world sensory and motor dynamics. In such cases, inference must rely on approximate methods, such as extended or unscented Kalman filters, particle filters, or more structured approximations like variational filtering [24, 37]. Incorporating these methods would allow our SLDS framework to capture more realistic sensorimotor transformations and state transitions. However, this introduces an additional challenge of performing approximately optimal control in these non-linear settings. A potential solution would be to use the iLQG method [38–40] which works by iteratively linearising the non-linear system via a Taylor expansion along some nominal trajectory (given some initial estimate of the states) and then uses this to recompute a better trajectory. A limitation is that the iLQG method assumes a fixed, known non-linear system, unlike the mixture of systems considered within the present SLDS framework.

Another critical extension concerns parameter learning. Currently, the model assumes fixed parameters (e.g., observation noise, dynamics), but the model could be extended by inferring them directly from the data. Approaches such as expectation-maximization (EM) or online Bayesian methods would allow the participant to adapt its internal models through experience [29, 41]. When combined with causal model inference, this would allow for the modelling of a more adaptive and flexible participant, capable of calibrating its internal dynamics alongside its understanding of causal structure.

Importantly, our current formulation assumes that the participant does not control the causal structure itself — it passively infers it. Future models could investigate how participants might intervene to test or alter their causal hypotheses, moving towards active causal inference frameworks.

Another potential direction for future research is to move from a finite set of discrete causal models to more flexible, non-parametric representations, such as those used in the COIN model [29], where causal structures can be inferred, updated, or created as needed. This would enable the system to both switch between existing models, recalibrate models (update parameters) and expand the model space to help learn particularly novel dynamics. A key question is how to expand the approach we consider here to this non-parametric formulation which is amenable to real-time control.

Finally, while the current model is computational, understanding how such a system could be implemented in the brain is a critical open question. Proposals have been made that probabilistic population coding (PPC) can implement Kalman filtering [42] or support static causal inference [43]. However, unifying these approaches into a biologically plausible neural circuit remains unresolved (although see [44] for a recent review on multisensory coding relevant to self-motion). Rideaux et al [45] demonstrated an artificial network can be trained to perform multisensory causal inference for visual-vestibular systems. Inspection of the network units, revealed a combination of congruent and opposite neurons, as found in neurophysiological studies [46]. Yet, this work does not address the changing sensory inputs over time or closed loop control, leaving open the question on how the type of systems studied here could be explained in terms of neural mechanisms.

## Methods

### Data

We extracted the mean data from [7, 19] using WebPlotDigitizer [47]. This software allows for the data to be manually extracted from plots using defined axes. As such the presented data may differ slightly (due to motor error) from the data shown in the original papers. The extracted data and code used to generate the results can be found at https://gitlab.socsci.ru.nl/sensorimotorlab/causal_inference_with_control

### Modeling framework

Our goal is to develop a modelling framework that explains control-based tasks involving causal inference. In such tasks, the participant is unsure about the true model that generated their sensory observations. This entails a dual problem; the participant must both infer the model and generate appropriate motor commands to complete the task [14, 25]. Importantly, we want our framework to be applicable to biological sensorimotor systems, requiring it can account for the different delays of sensory systems [48] as well as for signal-dependent noise in both the motor and sensory systems [12, 13, 27], while performing causal inference. The full pseudo code for the GPB2 algorithm is given in Algorithm 1. Before we describe it in detail, we first define the system on which inference is performed then specify how control signals are generated.

### System

We consider a finite set of *M* potential causal models, denoted by 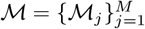, between which the participant must adjudicate. At each time step *t* ∈ ℕ, the active model is indexed by a random variable *z*_*t*_. The evolution of this model index follows a Markov process governed by a transition probability matrix *P* ∈ ℝ^*M* ×*M*^, such that

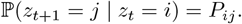

We assume that the external environment is governed by a latent dynamical system with a fixed (but unobserved) state 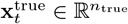. The system produces observable outputs **y**_*t*_ ∈ ℝ^*p*^ and responds to control signals **u**_*t*_ ∈ ℝ^*m*^ generated by the participant. However, each internal model ℳ_*j*_ maintained by the participant is not required to share the same state dimensionality. Instead, each model defines an internal latent state 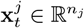, where *n*_*j*_ may vary across models. Each model ℳ_*j*_ thus defines a generative process over its own internal state and observations, with the following linear Gaussian dynamics that incorporate both additive and multiplicative noise components [13, 27]

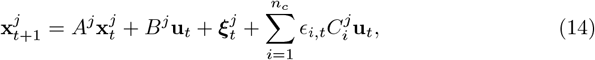

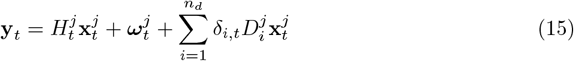

in which, 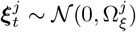 and 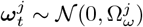 are both zero-mean additive Gaussian noise reflecting process noise and observation noise, respectively. The scalar random variables *ϵ*_*i,t*_, *δ*_*i,t*_ ~ 𝒩 (0, 1) are i.i.d. and model multiplicative noise. The matrices 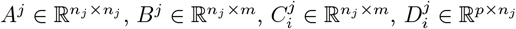, and 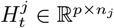 define the dynamics and observation mapping of model ℳ_*j*_.

This formulation allows internal models to differ in complexity and structure from the environment. For example, a model may represent a stationary object using only a position variable (*n*_*j*_ = 1) or equally by another with position and a latent zero velocity (*n*_*j*_ = 2). In such cases, the observation mappings 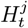 (or more generally, functions 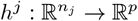) reconcile the internal representation with the observed output space.

Thus, even though models may differ regarding the dimensionality or structure of the underlying state, they can still be compared in a shared observation space. For simplicity in our simulations considered here we force equal dimensionality but providing 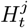 is appropriately designed, the algorithm operates correctly regardless.

### Estimation & control: Sub-models

The system defined if *j* is given and known (i.e., the model is known) has been used widely to model reaching movements [12, 13, 48, 48], grasping [23] and object manipulation [49]. The presence of state- and control-dependent noise renders the optimal policy intractable (it violates the certainty equivalence principle [14]). However, by assuming a linear prediction-based estimator, it is possible to derive an approximately optimal policy for this class of systems [13]. More specifically, the respective algorithm involves combing a linear state estimator paired with a linear feedback controller to compute a motor command. In this original work [13], two forms of the estimator are provided: an ‘adaptive’ and a ‘non-adaptive estimator’. The non-adaptive estimator is found by computing the optimal Kalman gain by averaging over potential states and control outputs, whereas the adaptive estimator is allowed to be a function of the state estimate 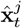 and control signal **u**_*t*_. Given we intent to merge multiple models, and thus the error dynamics in our system may be substantially different, we opt for the ‘adaptive’ estimator. This yields a predictive estimator for each model ℳ_*j*_ which gives an estimate 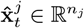 of the latent state (under the models assumptions). The estimator evolves according to the following update equations:

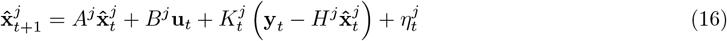

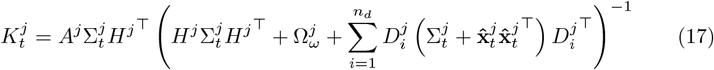

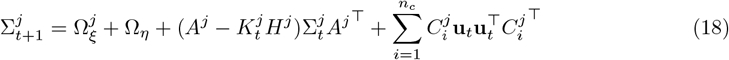

Here, 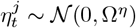 represents internal estimator noise, and 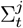 is the estimated state covariance. The gain matrix 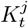 generalizes the Kalman gain to account for multiplicative noise in the observations via the additive term involving 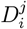, while the process covariance update includes analogous control-dependent noise through 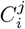.

The control output is a linear function of the state estimate, given by,

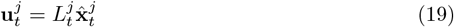

where L is the feedback gain matrix. We refer to the original paper [13] for the recursive equations necessary to compute this L matrix.

### Estimation : over model

During causal inference we cannot treat the latent model *z*_*t*_ as a known variable, it must be inferred, along with the state. To formalize this, we assume that a participant has a set of models which known parameters. More formally, let 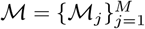 represent the set of candidate models, each associated with latent states 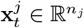 and corresponding system matrices.

### State propagation

At each time step *t*, the participant begins with a predictive state estimate 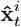 and covariance 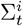 for each model according to the predictive estimator dynamics given in equations (16)–(18). Given these estimates 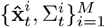 the joint one-step-ahead predictive estimates for all mode transitions (*i, j*) are computed by propagating 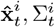 through the estimator associated with model ℳ_*j*_:

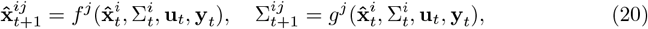

where *f* ^*j*^ and *g*^*j*^ correspond to the predictive state and covariance update defined by equations (16) and (18) respectively.

### Update

The likelihood of the current observation **y**_*t*_ conditioned on the transition (*z*_*t*_ = *j, z*_*t*−1_ = *i*) is given by the Gaussian density

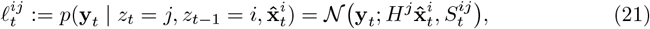

with covariance

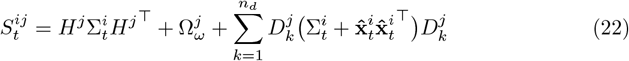

which accounts for additive and multiplicative observation noise under model ℳ_*j*_.

We can use this to perform a joint posterior probability update of mode transitions via Bayes’ rule:

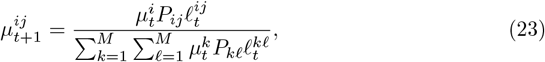

Where 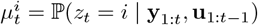.

### Mixture collapsing

Marginalizing over previous modes yields the updated mode probabilities at time *t* + 1:

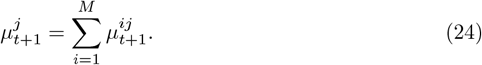

The predictive state estimates and covariances for mode *j* are obtained by collapsing the mixture of transitions:

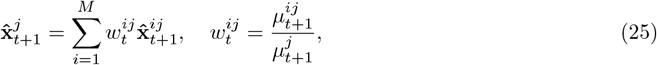

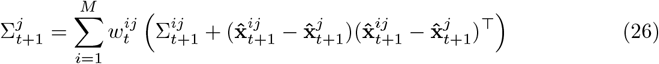

This mixture then serves as the input to the next step of the iteration.

### Control : over models

Previous causal inference work has modelled the response behaviour using the framework of Bayesian decision making [2, 8], where a loss function is given and a response is provided which minimizes the expected value of the loss. For example, initial work on perceptual causal inference [8] assumed a quadratic loss, and so the optimal estimate is the mean of the posterior distribution over the relevant variable.

Unfortunately, extending this to the case where continuous sensorimotor control is needed is far from trivial. In continuous control, a participant must try to accomplish the task in the face of model uncertainty. The optimal policy is generally intractable in this case, except for very select systems (e.g., LQG). As such we opt for a heuristic approach similar to the Bayesian averaging approach. That is, the participant forms a belief-weighted control signal by averaging over models using the current mode probabilities 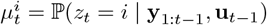:

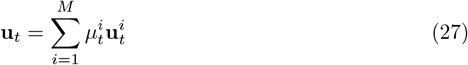

#### Algorithm 1

GPB2 with mode-weighted control

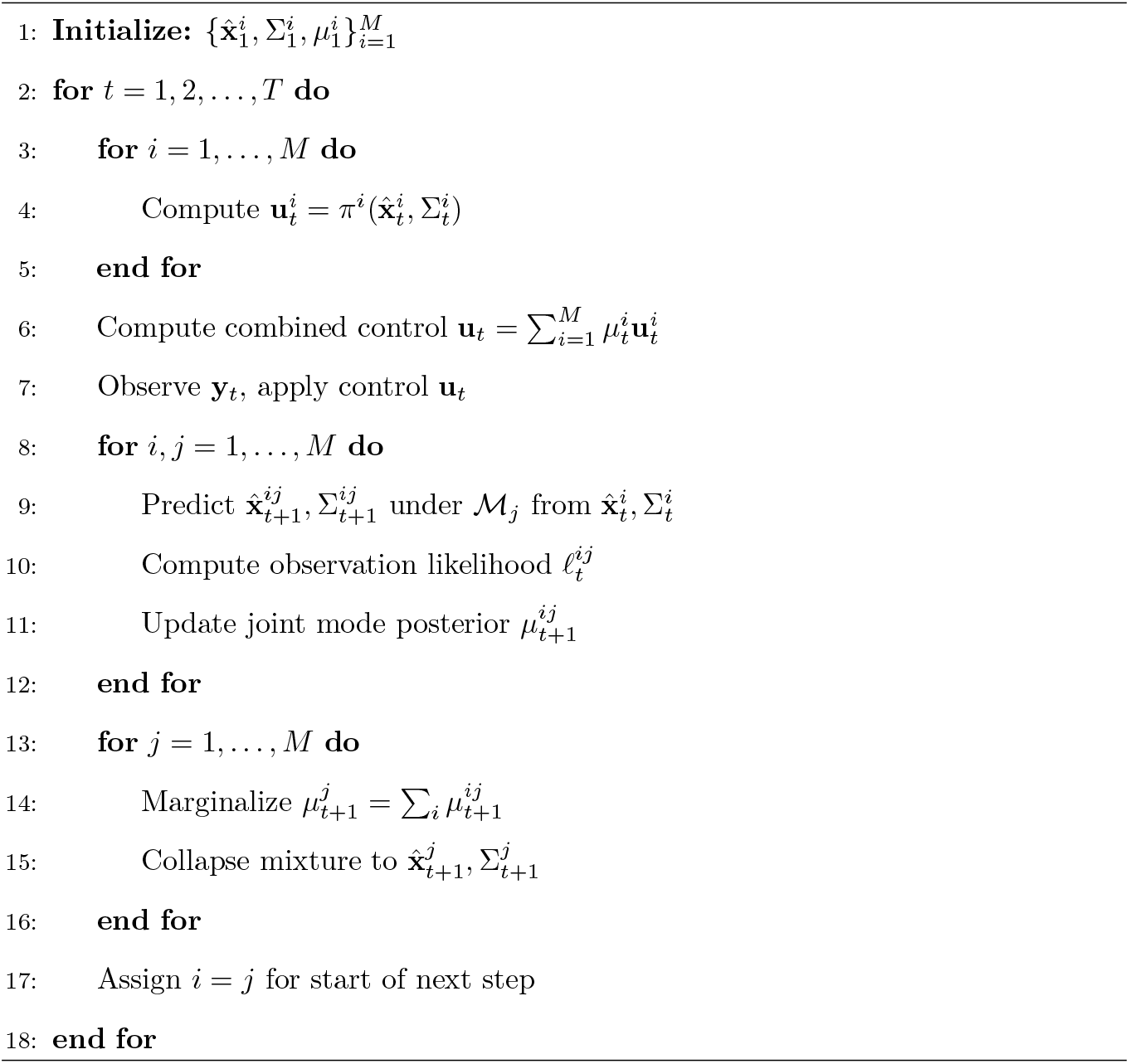

### Simulations

We simulated all the task by building on GYMNAX [50]. This leverages JAX [51] to allow for efficient computation. We also implemented our main algorithm in JAX for similar reasons. We obtained synthetic data for plotting by Monte Carlo simulations of 10000 trials.

In order to demonstrate applications of our algorithm we selected two examples from the causal inference literature (focusing specifically on visual-vestibular processing), a passive dynamic task [7, 17] in which the participant must determine whether an object moved during concurrent self-motion (in some heading direction) as well as distinguish heading direction from straight ahead. The second example was a similar but active dynamic task [19]. The participant was required to intercept a target which could either move or remain stationary. The participant only saw the target briefly and then had to navigate and stop to intercept it using ground based optic flow. The background flow used a fixed lifetime stimulus to prevent absolute position cues. Below we provide more details on each of the simulations, the necessary parameters for the simulations are given in Table 1.

**Table 1.** Parameters for the different tasks. Task subheadings indicate parameters are used in the task simulation, participant subheadings indicate parameters used by the participant. In case of overlap (e.g both use a parameters) they are set to equal.

| Parameter | Symbol | Value / Description |
| --- | --- | --- |
| **Dokka parameters** |  |  |
| *Task:* |  |  |
| Time step | $\Delta t$ | 0.01 s |
| Object speeds | $v_{\text{obj}}$ | $\{-5.35, \dots, 0, \dots, 5.35\}$ m/s |
| Velocity pulse width | $\tau$ | 1/6 m/s |
| Time | T | 1 (s) |
| *participant:* |  |  |
| Velocity decay | $\alpha$ | 0.99 |
| Visual background velocity noise SD | $\sigma_b$ | 0.15 m/s |
| Visual object velocity noise SD | $\sigma_o$ | 0.35 m/s |
| Acceleration noise SD | $\sigma_a$ | 0.2 m/s<sup>2</sup> |
| Body acceleration (process) | $\sigma_a^p$ | 2 m/s<sup>2</sup> |
| Object velocity (process) | $\sigma_o^p$ | 0.2 m/s |
| **Noel parameters** |  |  |
| *Task:* |  |  |
| Time step | $\Delta t$ | $\frac{1}{90}$ (s) |
| Observation visibility duration | $T_{\text{view}}$ | $\{7, 13\}$ steps |
| Maximum steps per episode | $T_{\text{max}}$ | 135 steps |
| Target speed | $v_{\text{init}}$ | $\{0, 0.03, 0.06, 0.10, 0.20, 0.40\}$ m/s |
| Target distance | $d$ | 2.0 m |
| *participant:* |  |  |
| Position noise SD (target $x$) | $\sigma_{\text{pos},x}$ | 0.005 m |
| Position noise SD (target $y$) | $\sigma_{\text{pos},y}$ | 0.009 m |
| Velocity noise SD | $\sigma_v$ | 0.01 m/s |
| Velocity prior SD | $\sigma_{\text{vel,prior}}$ | 0.5 m/s |
| **Switching Extension** |  |  |
| Unlimited observation visibility | $T_{\text{view}}$ | ∞ (no zeroing of target position) |
| Velocity switching interval | $T_{\text{switch}}$ | 30 steps |

### Dokka task: Object stationarity and heading direction

In this task, the task participant’s body follows a fixed minimum-jerk displacement trajectory over a duration *T* with time step Δ_*t*_, yielding a smooth position profile in both lateral (*x*) and depth (*y*) dimensions. The trajectory is defined such that the body moves from an initial position to a final position with a total displacement of 0.13 m in a particular heading angle (see [7] for more details).

During this self-motion an object (moving horizontally) is assigned a velocity profile constructed from a scaled Gaussian,

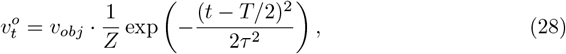

where *v*_*obj*_ denotes the peak speed (sampled from a predefined set of values), *τ* controls the width of the velocity pulse, and *Z* is a normalization constant ensuring 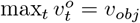. The object position 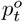 is obtained by discrete integration of 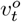.

We can define the full latent state of the body(^*b*^) and object (^*o*^) at time *t*

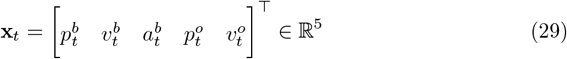

with independent instantiations for both *x* and *y* dimensions, resulting in a 10-D vector in total.

Sensory observations (see Eq 15) are generated as noisy measurements of the state, corrupted by additive Gaussian noise. Specifically, we simulate three observations per spatial dimension:

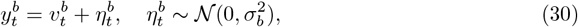

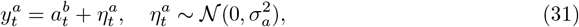

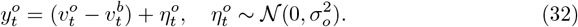

This abstracts the early visual processing stages out and assumes that optic flow is parsed into global (background) and local (object-relative) motion components. This gives three sensory observations, a visual estimate of body velocity via background optic flow: 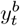, a vestibular estimate of body acceleration: 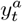, and a visual measurement of object velocity via local motion around the object: 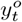. This yields the total measurement vector,

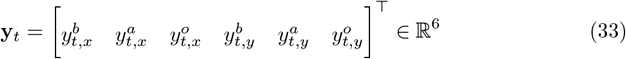

In the visual-only condition, vestibular acceleration signals are removed from the observation vector by setting the corresponding measurement to zero. The resulting observation vector at each time step is

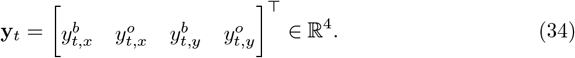

### Dokka: Participant models

From the standpoint of the participant, we assume that the body can be represented as a moving point mass [13] governed by Newtonian dynamics with acceleration noise (acceleration fluctuates randomly fluctuates over time). We combine this model with a simple target velocity model [52]. Thus, the dynamics of the body in discrete time (for each dimension, below is the x dimension) as,

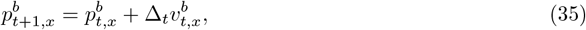

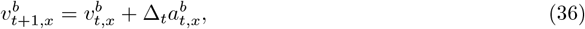

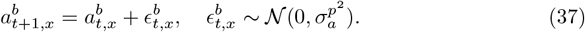

We concatenate motion in both lateral (*x*) and depth (*y*) directions by stacking the dynamics matrices into a block-diagonal form, yielding a full state-space representation for body motion.

Of the object, we assume a linear autoregressive velocity model, as commonly used in motion perception literature [52]:

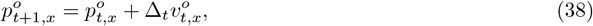

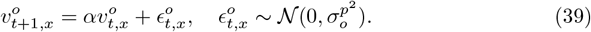

We determined the bias in terms of the heading percept (via the models estimated body position at the end) and the proportion of times the model indicated the object did not move in a whole trial (no probability of motion 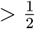).

### Noel task: Path integration interception

For a full understanding, we first detail their task. In their experiment, participants were shown a target in front of them for a certain amount of time (75 or 150 ms), after which they proceeded to try and intercept the target. This was done by moving a joystick which controlled the angle and speed of their motion (e.g. they could move towards the target at a speed they controlled). In 25% of trials the target stayed stationary and in the remaining 75 % moved at a constant speed (see Table 1). We simulate the task using a latent dynamical system governing the motion of a planar target (*T*^*^) and a body (B). The true latent state at time *t* is given by

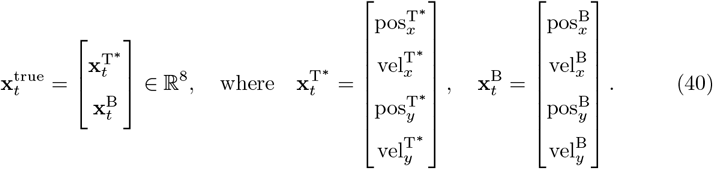

The target moves at a constant (but sampled) velocity in a straight line. The body is controlled via acceleration inputs **u**_*t*_ ∈ ℝ^2^, corresponding to lateral and depth accelerations. The system evolves in continuous time with discretization step size Δ*t* = 1*/*90. The dynamics are given by (with ⊗ the Kronecker product):

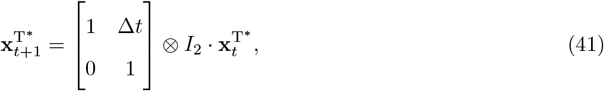

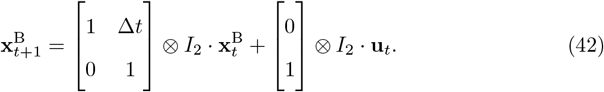

At each time step, a noisy observation **y**_*t*_ ∈ ℝ^4^ is produced, reflecting the target position and body velocity:

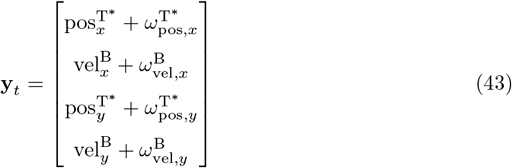

where each noise term is independently sampled from a zero-mean Gaussian distribution:

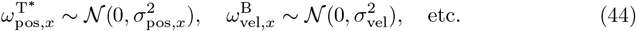

Visibility of the target is restricted to a fixed time window *T*_view_, after which the target position components of **y**_*t*_ are set to zero, hence

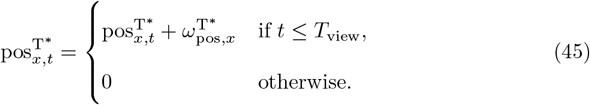

The initial target distance and speed were set to match the original experiment [19] (see Table 1).

### Noel: Participant models

Noel et al [19], employed a few simplifications to derive a tractable participant model. For estimation they assumed that depth is known perfectly so the participant only needs to infer the velocity of the target to decide whether it is moving or not. For control the authors assume that after the target disappears, the participant plans an open-loop policy to intercept the target by moving in a straight line (constant velocity) at a given angle *θ*^*^ and interception time (t). To this end, the authors assume the participant’s estimation of self-location becomes worse as a function of distance. In contrast, in our formulation, both depth and lateral position are estimated and control is implemented in a closed-loop manner.

The participant maintains two competing internal models to explain and predict the observed sensory data: a *motion model* that assumes the target moves with constant velocity, and a *stationary model* that assumes the target is fixed in position. Both models share the same internal state representation but differ on in the assumed dynamics of the target component.

Let the internal latent state for each model be defined as

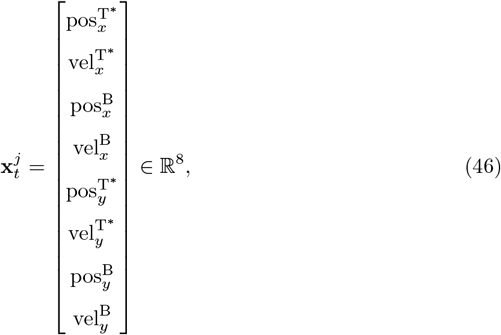

where superscripts T^*^ and B refer to the target and body respectively, and subscripts *x, y* denote lateral and depth components. The participant controls the body acceleration via a 2D input **u**_*t*_ ∈ ℝ^2^, which directly modifies body velocity.

### Motion model

This model assumes that the target moves with constant velocity. The internal dynamics follow a linear Gaussian system:

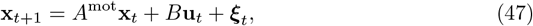

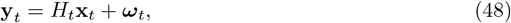

where ***ξ***_*t*_ ~ 𝒩 (0, Ω_*ξ*_), ***ω***_*t*_ ~ 𝒩 (0, Ω_*ω*_), and there is no multiplicative noise (i.e., *C* = *D* = 0). The dynamics matrix *A*^mot^ ∈ ℝ^8*×*8^ assumes constant velocity for both target and body, while *B* ∈ ℝ^8*×*2^ maps control inputs to the body velocity:

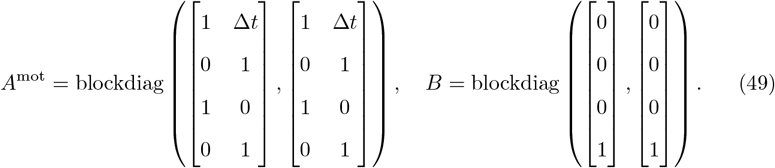

The observation model *H*_*t*_ switches at time *T*_view_. Before this time, the participant observes both target position and body velocity; afterwards, target position is unobservable:

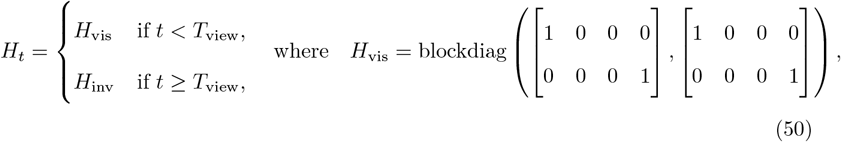

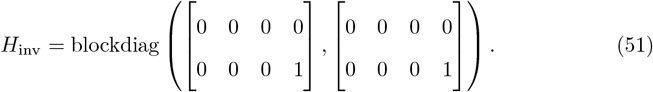

### Stationary model

The stationary model is structurally identical to the motion model but assumes that the target does not move over time. This is achieved by altering the target dynamics block in *A* to remove velocity propagation:

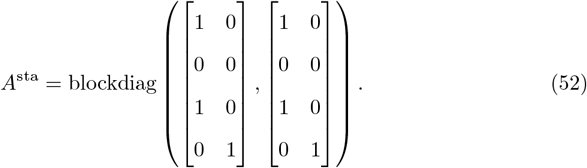

The dynamics matrix *A*^*sta*^ ∈ ℝ^8^ produces a body state that evolves according to Newtonian dynamics, but a target position that remains stationary. All other matrices are as defined in the motion model.

### Model Initialization and Cost

Each model starts with a prior **x**_1_ ∈ ℝ^8^ centred on a target located directly ahead, and a body at the origin:

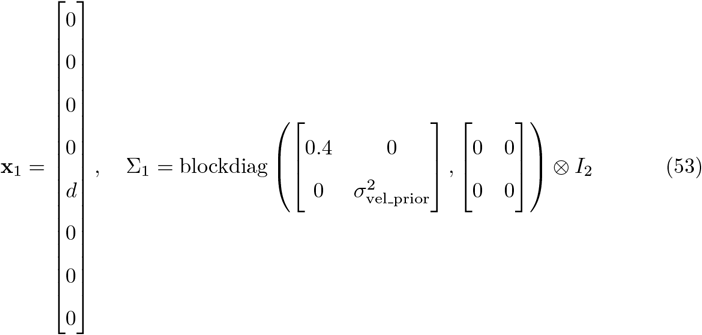

Each model defines a final-time cost encouraging the body to be aligned with the target’s final position:

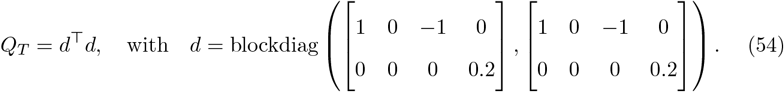

This cost encourages spatial alignment (position matching) and penalizes terminal velocity errors with lower weight. This matches the task structure where participant had to stop to indicate the end of a trial.

